# Mapping the Dark Side: Visual Selectivity of Default Network Deactivations

**DOI:** 10.1101/292524

**Authors:** Tomas Knapen, Daan van Es, Martijn Barendregt

**Affiliations:** Vrije Universiteit Amsterdam, Cognitive and Applied Psychology, van der Boechorststraat 1081BT Amsterdam, The Netherlands; Spinoza Centre for Neuroimaging, Meibergdreef 75, 1105BK Amsterdam, The Netherlands

## Abstract

The default network (DN) activates for internally referenced mental states, whereas it deactivates in response to sensory stimulation. We show that BOLD signal decreases in the DN are tuned to the spatial location of visual stimuli, allowing us to decode the location of a visual stimulus from DN regions. Our results indicate that the DN represents sensory information, and suggest it may utilize this reference frame for higher-level cognition.

---

The default network (DN) deactivates when participants receive sensory stimulation and focus externally to perform a task, and activates during mind-wandering^1,2^. These activations and deactivations in the DN are important in psychological disorders and alterations in cognitive state^3,4^, and are inversely correlated with responses in the multiple demand network of frontal and parietal regions^5^. Recent findings indicate that signals in the DN carry visual memory information^6,7^, but the functional role of DN deactivations remain unclear. We investigated the visual response properties of the DN by performing a visual mapping fMRI experiment at ultra-high field (7 Tesla). In our experiment a bar-shaped stimulus systematically traversed the visual field in different directions. Participants performed a two-alternative forced-choice color discrimination task on the stimulus (Figure 1a) titrated to be equally demanding throughout the visual field (Figure 1b). Across participants, there was no difference in task performance (F(2,10) = 1.347, p = .304, η^2^ = .198) or reaction time (F(2,10) = 0.121, p = .887, η^2^ = .003) for different locations of the visual stimulus. In addition, we verified that the bar stimulus did not induce eye movements, as the variability in gaze direction did not depend on the direction of the visual stimulus (See methods, F(1,10) = 0.222, p = .657, η^2^ = .018), for all bar stimulus eccentricities. These results show that task difficulty and attentional load did not change as a function of stimulus location and rule out the possibility of confounding task-related signals with spatially selective signals.

**Figure 1:**
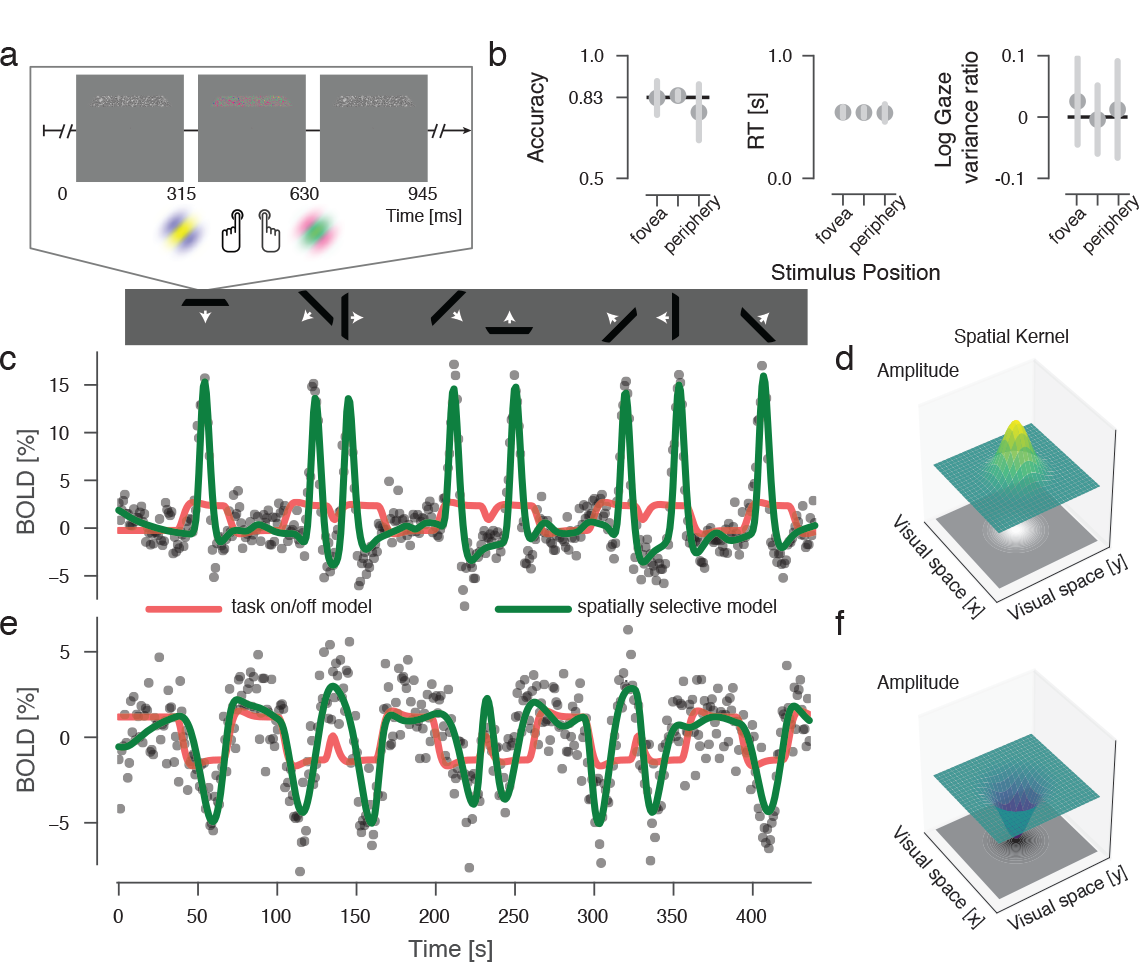
Behavior and single-voxel responses. **a.** Stimulus and task design. **b.** Behavioral results. There was no significant difference between accuracy, reaction time and gaze variability as a function of the bar stimulus locations. **c-d**. BOLD signal time-course from a single V1 voxel. This response is best explained by a spatially selective model (green line, implemented as spatially localized Gaussian kernel with a positive amplitude) than by a simple task on/ off model (red line). **e-f**. BOLD signal time-course of a single angular gyrus voxel. Contrary to V1 responses, BOLD signals decreased when a stimulus was presented within the voxel’s pRF. This response is best explained by a spatially selective model (a Gaussian kernel with a negative amplitude, green line), rather than by the task on-off model.

Turning to the recorded BOLD responses, Figure 1c shows the time-course of visual stimulation and the resulting BOLD fluctuations from a single cortical location in primary visual cortex. As expected in this brain region, the structured visual input caused BOLD increases only when the stimulus was present within a circumscribed region of the visual field (see Figure 1d), mathematically modeled as a population receptive field (pRF)^8^. This spatially selective model shows an excellent approximation to the BOLD signal time-courses (across-runs average cross-validated (CV) *R*^2^ =0.82), compared to a model that has no visual field selectivity and only encodes stimulus/task presence (CV *R*^2^ =0.10). We then applied identical analyses to BOLD signals arising in the DN, from the same recordings. Figure 1e shows an example signal time-course from the lateral parietal angular gyrus and illustrates the strong BOLD signal decreases observed whenever a stimulus was presented.

Our spatially non-selective model fit the deactivations of this cortical location during stimulus presence (CV *R*^2^ =0.15), confirming that it is not only anatomically but also functionally part of the DN. However, the strongest deactivations are not captured by this spatially non-selective model. We therefore fit a spatially selective model that implements a negative population receptive field (Figure 1f). This model captures the specific temporal structure of deactivations, showing much better cross-validated prediction performance than the spatially non-selective model (CV *R*^2^ =0.63). These results indicate that DN negative BOLD responses encode visual location, which can be computationally formalised as a spatial pRF of negative amplitude. This pattern of results was typical for recordings from the DN, and we selected only voxels where the spatially selective model outperformed the non-selective model for further analysis. Figure 2 shows inflated and flattened cortical surface depictions of the predictive power of our spatially selective model, colored based on whether they result from activations (orange colormap) or deactivations (blue colormap). The spatial distribution of activations across the cortical surface confirms earlier findings of visual selectivity in parietal and frontal cortex^9^. The spatial distribution of deactivations on the other hand corresponds well with the locations of DN nodes in parietal and frontal brain regions as found in resting-state experiments which used large populations of participants^10,11^. Across participants, there was some variability in the exact locations of the clusters of negative pRFs (supplementary Figure 2). These individual-level specificities of deactivations are reminiscent of recent findings showing fine-grained connectivity between DN nodes^12^.

**Figure 2:**
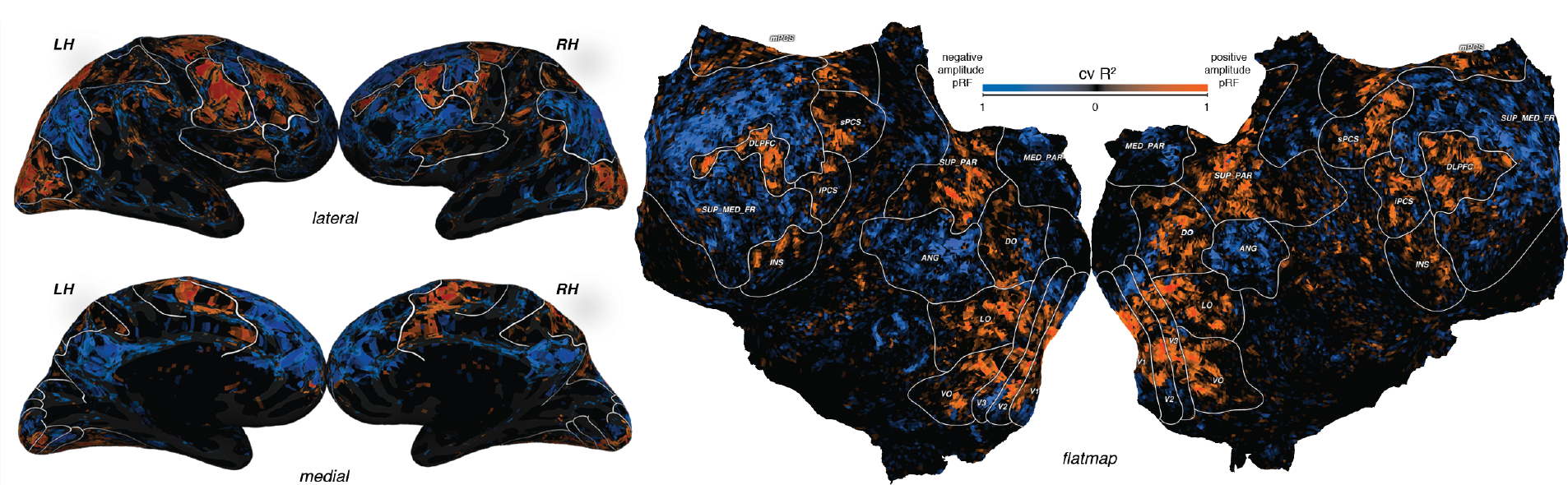
Inflated and flattened depictions of positive (red) and negative (blue) pRFs across the cortical surface of an example subject. VO: ventral occipital cortex encompassing visual field maps V4 and VO1/2; LO: lateral occipital cortex encompassing visual field maps LO1/2 and TO1/2; DO: dorsal occipital cortex encompassing visual field maps V3AB, V7 and IPS0/1; SUP_PAR: superior parietal lobe, encompassing visual field maps IPS2/3/4/5; sPCS: superior precentral sulcus; iPCS: inferior precentral sulcus; mPCS: medial precentral sulcus; DLPFC: dorsolateral prefrontal cortex; INS: insula; ANG: Angular Gyrus; MED_PAR: medial parietal lobe; SUP_MED_FR: superior & medial cortex. Supplementary figure 2 provides these flat map depictions of all subjects, as well as maps of CV R^2^ of the task on/off model, and the difference in CV R^2^ between selective and non-selective model.

We then asked if the pattern of deactivations within separate DN nodes encode the location of a stimulus in the visual field (Figure 3). Specifically, we used pRFs parameter estimates from a given cortical region as an explicit encoding model. We then used a recently developed Bayesian decoding algorithm^13^ to calculate the most probable stimulus pattern given the structure of BOLD responses across voxels in a test dataset, separately for every TR (Figure 3a). Performing this analysis results in movies of inferred stimulus representations for the entire experiment (Figure 3b-c). We quantified the fidelity of a brain region’s representation of visual field location of a stimulus by means of the correlation between the actual stimulus location and the inferred stimulus location. Figure 3d shows that stimulus position can be decoded from known parietal and frontal retinotopic areas (two-tailed t-test on median across CV folds, sPCS: t(5)=5.48, p=0.003, iPCS: t(5)=4.13, p=0.009, SUP_PAR: t(5)=6.86, p=0.001), reflecting their visuospatial organization. The same analysis, performed on DN nodes, found significant correlations between actual and decoded stimulus position (ANG: t(5)=4.80, p=0.005, MED_PAR: t(5)=3.264, p=0.022, SUP_MED_FR: t(5)=4.13, p=0.009, Figure 3e). In sum, we find significant decoding performance for DN nodes with a fidelity similar to that of known retinotopic regions in higher-level visual cortex (Figure 3f).

**Figure 3:**
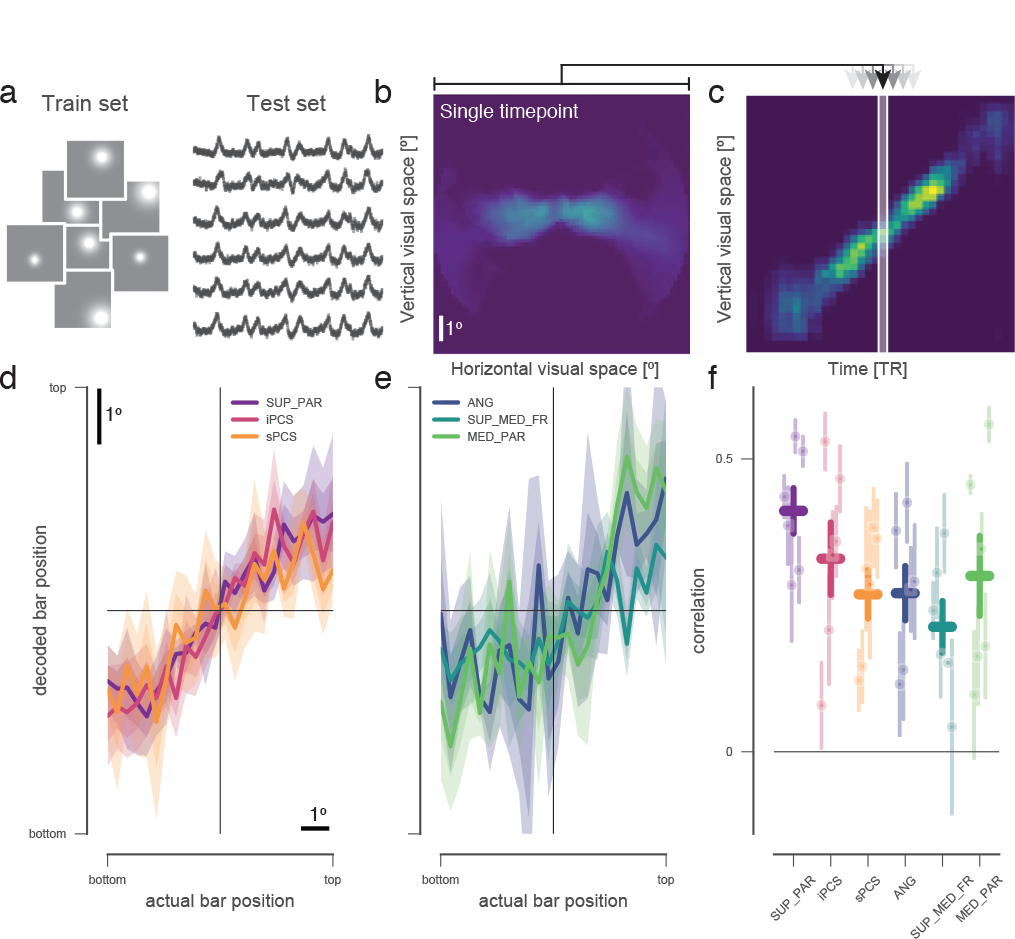
Decoding of visual location. Using a forward model, we decoded the location of our visual stimulus in a cross-validated fashion. **a.** pRF fits from a training set were combined with voxel time-series from an independent test set. **b.** We computed the most likely spatial stimulation based on the pattern of voxel responses on every time point^13^, example from visual area V2. **c.** These images were then rotated relative to the bar’s direction, averaged first across stimulus directions, and then across the direction perpendicular to the stimulus. The center of mass of the resulting spatial distribution for each timepoint was taken as the decoded bar position (after shifting by the haemodynamic delay of the BOLD response). **d.** Known high-level retinotopic maps in parietal and frontal cortex show a strong correlation between actual and decoded stimulus position. **e.** DN regions allow decoding of visual stimulus location based on their BOLD signal decreases. **f.** Decoding results across regions and participants. Error bars represent the standard error of the mean across participants and CV folds. Supplementary figure 3 shows results of this analysis for low-level visual regions.

These results indicate that DN areas encode relatively detailed visual location information by means of their deactivations. The DN is thought to constitute one extreme of a gradient leading from primary sensory and motor regions to transmodal association cortex^14,15^. What would be the use of the brain representing visual location in these high-level regions that integrate across sensory modalities? We offer two potential and non-exclusive computational roles for this neural signature. First, the well-known signal increases of these DN regions in memory^16^ and social semantic information processing^17^ indicate that the DN activates for computations that emphasize internally instead of externally oriented information processing. The balance between the multiple demand and default networks could serve to tune processing either outwards or inwards, i.e. towards incident sensory information or, conversely, towards memory-based and/or ego-referenced processing. Visual space could serve as a shared reference frame for this interaction between networks. Second, there is abundant evidence from sensory processing that the interplay between activations and deactivations can serve to decorrelate neural responses to input patterns^18^ and increase processing efficiency by means of predictive coding^19^. Especially in the matching of multiple input patterns there is inherent computational benefit to representing what is not there, as opposed to solely representing what is there^20^. The DN is ideally suited to perform this type of matching operation on the level of transmodal association^15^ and memory^16^. Our results suggest that the spatial arrangement of sensory inputs may be frugally inherited by the DN to support higher-level cognition.

## Acknowledgments

We would like to thank Serge Dumoulin, Alex Huth, Martin Szinte, and Gilles de Hollander for fruitful discussions and comments on an earlier version of this manuscript, and Marco Aqil for software engineering. TK was supported by NWO-CAS grant 012.200.012 and ABMP grant 2015-7. The authors declare no competing financial interests.

## Methods

### MR procedures

Functional MRI were acquired using a 7T Philips (Best, NL) Achieva scanner at the Spinoza Centre, Amsterdam, the Netherlands, with a dual-channel transmit and 32 channel receive coil (Nova Medical, Wilmington, MA, USA). A Gyrotools (Zürich, CH) simultaneous multi-slice selection (factor 3) echo-planar imaging sequence was used, with a resolution of 2 mm isotropic, a repetition time (TR) of 0.945s, an echo time (TE) of 29 ms and a flip angle of 56°. K-space data were exported and reconstructed to image space offline. A T1 weighted anatomical scan (1 mm isotropic, TR 8000 ms, TE of 3.73 ms, flip angle of 8°) was acquired on a Philips 3T Achieva equipped with a 32-channel head coil. Cortical surfaces were automatically segmented using freesurfer 5.3, followed by hand-editing of the white-gray matter boundary and pial surfaces. Six participants (1 female) were scanned during 1.5 hour sessions. All participants had normal or corrected-to-normal acuity and normal color vision. During scanning we performed eye tracking (Eyelink, SR Research, Osgoode, CA) at 1kHz, and used eye position information to verify accurate fixation of participants. Runs in which participants made saccades to the stimulus were removed from further analysis.

### Stimulus and Behavior

Traveling bar stimuli were used for retinotopic mapping according to a standard population receptive field imaging design^8^, see Figure 1. The experiment was created in python using pygaze and psychopy packages, and the code implementing the experiment is available at https://github.com/VU-Cog-Sci/PRF_experiment/tree/7T_exp. Stimuli were presented on a 32’’ full HD BOLDScreen (CRS, Cambridge, UK) at the end of the bore, and viewed through a front-silvered mirror. Bar stimuli were filled with 2000 drifting Gabors with a spatial frequency of 2 cycles per degree, random orientations, and temporal frequency of 4Hz. The bar containing the Gabors was 1.25 degrees of visual angle (dva) wide, and traveled through a 11 dva diameter circular aperture in 30.2s. Bar traversals and blanks were ordered as depicted in figure 1. The position of the bar was updated on every TR (32 steps), and the gabors were refreshed three times per TR, i.e. every 315 ms. Gabors were drawn in grayscale, except during the middle 315 ms of the TR, when the gabors were redrawn in cyan/magenta (CM) or blue/yellow (BY). The ratio between the number of CM and BY elements was under experimental control, and participants were required to perform 2 alternative forced-choice judgments on the color preponderance (more CM vs. more BY) on every TR. The difficulty of this task was titrated by means of three independent Quest staircases controlling the color preponderance at 3 different eccentricities of the bar stimulus, to ensure similar task difficulty across different eccentricities in the visual field.

### MR data analysis

All data analysis was performed using open-source software^21^.

Preprocessing was conducted using a nipype pipeline^22^ that performed B0 fieldmap-based spatial unwarping, motion correction (AFNI’s 3dvolreg), high-pass filtering (3rd order savitzky-golay filter, window width 120 s) and conversion to percent signal change referenced to the median signal value over time. All EPI runs were coregistered together to create a single functional space, and this space was coregistered to the subject’s T1 anatomical using FreeSurfer’s BBregister. We took the median across functional runs on a leave-one-out basis, to create up to 7 training runs. We fit pRF parameters for all gray-matter voxels in the functional space on each of these median runs while the left-out run was used as a test set in cross-validation procedures. Fitting of pRF parameters (x and y position, size, offset, amplitude and compressive nonlinearity power^23^) was performed using popeye^24^, with standard hemodynamic response function parameters. We verified that our results were not dependent on specific preprocessing parameters, by varying the temporal filtering characteristics, motion correction algorithm (FSL’s mcflirt vs 3volreg), and performing RETROICOR-based regression of cardiac/ respiration nuisance variables^25^. We furthermore performed pRF fits with a standard isotropic Gaussian model^8^, i.e. without the compressive spatial summation nonlinearity, showing similar results. Surface-based ROIs were defined based on per-subject retinotopic mapping (occipital, parietal and frontal visual areas) as well as on population-level templates of canonical default network regions^11^. Cortical surface analysis and visualization was performed using pycortex^26^. All code performing these analyses is available at https://github.com/tknapen/PRF_MB_7T.

### Decoding analysis

We adapted the decoding strategy conceived by Van Bergen et al^13^ for use in a scenario where the forward model feature space is visual location instead of orientation. We converted the continuous-valued pRF estimates to a discrete spatial representation of P by P pixel values. Thus, a weight matrix W of dimensions k by P^2^ described the spatial preferences of all voxels in a region of interest. All other aspects of the analysis were kept identical to the previous description^13^. Briefly, we fit a model describing the total covariance structure of voxels’ responses based on the overlap in their pRFs (WW^T^), as well as their noise variance and covariance. The inverse of the total fitted covariance matrix was used to estimate the most likely stimulus (i.e. pixel values) that evoked the test-set pattern of BOLD responses. All fitting was performed using L-BFGS-B gradient descent, implemented in scientific python. Varying the number of voxels used for the decoding analysis did not impact the pattern of decoding results, nor did varying the amount of feature space elements (pixels) used in the analysis. We did not add a spatial vicinity component in the forward model description as previous work has shown that this does not improve decoding performance^13^ and ensured that the fitted covariance matrix was positive-semidefinite. This analysis is implemented in an open-source python package, available at https://github.com/tknapen/decode_encode.

### Gaze analysis

We analyzed whether the bar induced eye movements by comparing gaze variability across orthogonal and parallel directions to bar movement^27^, as these eye movements can impact pRF estimates^28^. Gaze data was first cleaned by linearly interpolating blinks detected by the Eyelink software. Drift correction was performed by removing the median gaze position within each bar pass. Bar-passes in which less than 25% of the samples contained valid gaze recordings were excluded from analysis. We then computed the log-ratio of position variability (standard deviation) in the direction parallel compared to orthogonal of bar movement for each of three equally spaced bar position eccentricities. Positive values of this measure indicate greater eye position variability in the direction along compared to across bar movement.

